# A quantitative method for proteome reallocation using minimal regulatory interventions

**DOI:** 10.1101/733592

**Authors:** Lastiri-Pancardo Gustavo, J.S Mercado-Hernandez, Kim Juhyun, José I. Jiménez, Utrilla José

## Abstract

Engineering resource allocation in biological systems for synthetic biology applications is an ongoing challenge. Wild type organisms allocate abundant cellular resources for ensuring survival in changing environments, reducing the productivity of engineered functions. Here we present a novel approach for engineering the resource allocation of *Escherichia coli* by rationally modifying the transcriptional regulatory network of the bacterium. Our method (ReProMin) identifies the minimal set of genetic interventions that maximise the savings in cell resources that would normally be used to express non-essential genes. To this end we categorize Transcription Factors (TFs) according to the essentiality of the genes they regulate and we use available proteomic data to rank them based on its proteomic balance, defined as the net proteomic charge they release. Using a combinatorial approach, we design the removal of TFs that maximise the release of the proteomic charge and we validate the model predictions experimentally. Expression profiling of the resulting strain shows that our designed regulatory interventions are highly specific. We show that our resulting engineered strain containing only three mutations, theoretically releasing 0.5% of their proteome, has higher proteome budget and show increased production yield of a molecule of interest obtained from a recombinant metabolic pathway. This approach shows that combining whole-cell proteomic and regulatory data is an effective way of optimizing strains in a predictable way using conventional molecular methods.

**Importance:** Biological regulatory mechanisms are complex and occur in hierarchical layers such as transcription, translation and post-translational mechanisms. We foresee the use of regulatory mechanism as a control layer that will aid in the design of cellular phenotypes. Our ability to engineer biological systems will be dependent on the understanding of how cells sense and respond to their environment at a system level. Few studies have tackled this issue and none of them in a rational way. By developing a workflow of engineering resource allocation based on our current knowledge of *E. coli*’s regulatory network, we pursue the objective of minimizing cell proteome using a minimal genetic intervention principle. We developed a method to rationally design a set of genetic interventions that reduce the hedging proteome allocation. Using available datasets of a model bacterium we were able to reallocate parts of the unused proteome in laboratory conditions to the production of an engineered task. We show that we are able to reduce the unused proteome (theoretically 0.5%) with only three regulatory mutations designed in a rational way, which results in strains with increased capabilities for recombinant expression of pathways of interest.

**Highlights:**

- Proteome reduction with minimal genetic intervention as design principle
- Regulatory and proteomic data integration to identify transcription factor activated proteome
- Deletion of the TF combination that reduces the greater proteomic load
- Regulatory interventions are highly specific
- Designed strains show less burden, improved protein and violacein production

## Introduction

The removal of accessory non-essential functions is one of the strategies used to engineer microbial phenotypes. This approach relies on the assumption that cellular resources for gene expression are limited and, therefore, by removing unneeded genes in a certain environment, the cell is capable of allocating resources to other functions (e.g. expression of recombinant genes). These minimization approaches are mostly done by reducing genome size and gene number including performing random deletions^1,2^, however, the precise way in which the resource allocation takes place after the genetic intervention is not considered.

Organisms respond to the environment by cellular signalling encoded in regulatory networks^3^. The intricacy of the lifestyle of an organism is generally translated into signalling complexity^4^. Biological regulatory networks are robust^5^ and evolvable^6^ to cope with environmental and lifestyle perturbations, however this robustness involves intrinsic trade-offs, such as resource allocation strategies. It has been shown that cellular states are naturally “primed” for typical upcoming changes. Bacteria anticipate to fluctuations in the environment^7,8^, draining resources from functions that are mostly performed in relatively stable conditions. The expression of anticipation functions, also called hedging functions, is encoded in the regulatory network and it has a proteomic cost^9^. Genome scale models along with experimental data sets enable the calculation of the minimal theoretical proteome needed to sustain growth in a defined condition^10^. Therefore, comparing minimal theoretical proteomes with measured proteomes reveal the costs of the hedging proteome allocation. Proteome econometric approaches can facilitate the engineering of cellular states or phenotypes aimed at displaying an engineered function. Recent studies have focused in the host-construct interactions for increasing predictability of synthetic constructs^11–13^. In addition to these approaches, the rational design of the host used for expression following econometric models can be adopted to improve the performance of synthetic constructs, including production phenotypes for molecules of added value. Among other benefits, streamlined organisms obtained this way are less likely to develop undesired emerging behaviours^11^.

In this work we develop a new top-down cell engineering strategy for *Escherichia coli* using the Transcriptional Regulatory Network (TRN) as a control layer for proteome allocation. By combining high-throughput proteomic information, regulatory network interactions and gene essentiality observations, we develop a method capable of finding the minimal set of genetic interventions required to divert the resources invested in superfluous hedging into increased biosynthetic potential. The resulting strain exhibits an increased availability of cellular resources to express engineered functions.

## Results

### Combining gene essentiality and TRN analysis identifies dispensable TFs for proteome reduction in a defined condition

The genome scale model of Metabolism and gene Expression (ME model) computes the minimal theoretical proteome and allows calculating the cost of expressing hedging functions. It can be used to simulate different scenarios of expression of the hedging proteome (as unused protein fraction coefficient, see methods)^14^. These simulations allow us to calculate the costs and the benefits of different interventions, e.g. by modulating the expression of the hedging proteome, expressed in terms of growth, the size of both essential and recombinant proteome sectors (Supp. fig. 1).

**Figure 1.**
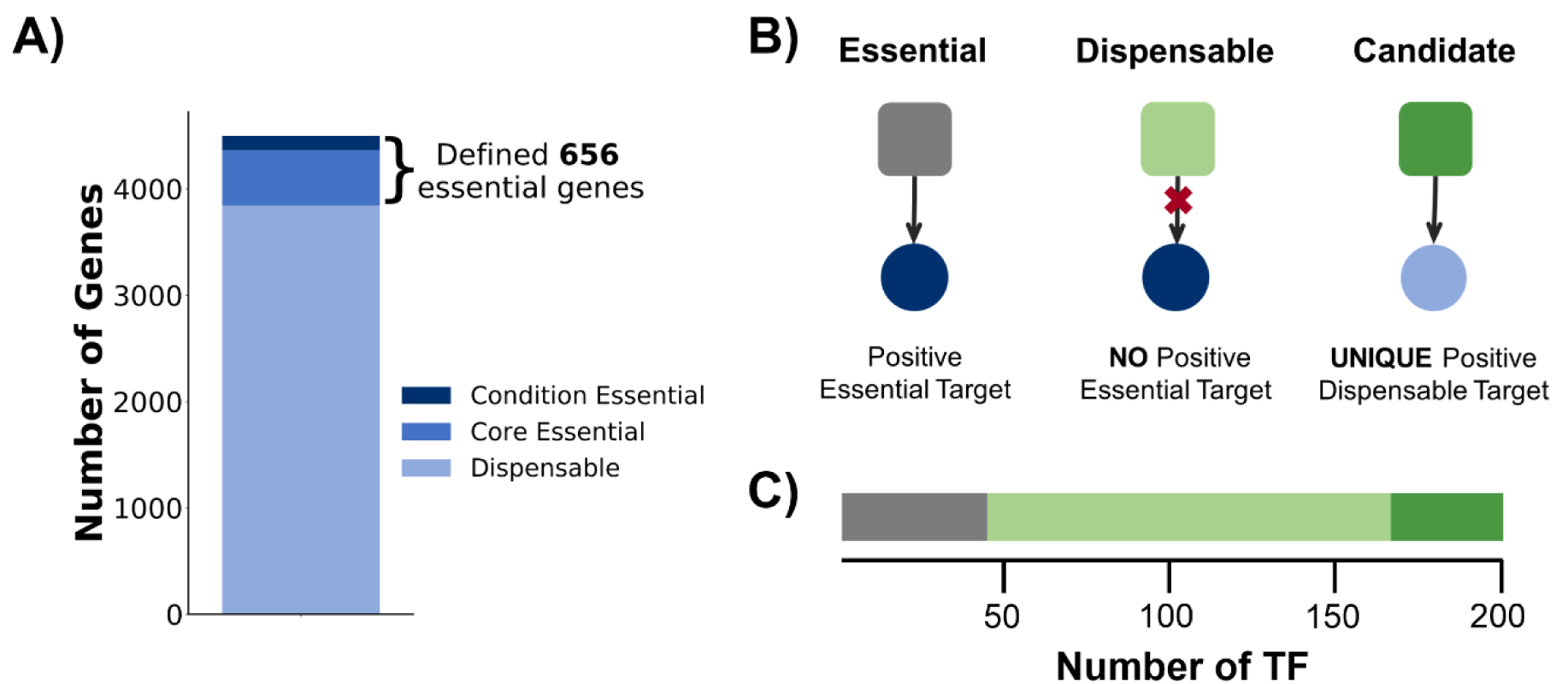
Gene essentiality and transcriptional regulatory network analysis in the pre-defined condition. **A)** Essentiality profile of *E. coli* genome under selected growth condition. The essential fraction of the genome is divided into core (always needed) and conditional (M9-glucose needed). **B)** Graphical representation of the sub-network of interactions considered for the classification of the TFs; grey squares represent essential TFs, light green squares dispensable TFs, green squares candidate TFs, dark blue circles essential genes, light blue circles dispensable genes and arrows positive interactions. **C)** Essentiality profile of the TFs contained in RegulonDB. Total *E. coli* genes according to Ecocyc^16^.

We build on ME-models to design strains containing the minimal genetic interventions that reduce the greatest amount of proteomic resources not required to grow in a specific condition. Our method uses Transcription Factors (TFs) as the key dials controlling the allocation of the hedging proteome in a pre-defined specific environment. We begin by establishing batch growth in minimal media (M9) supplemented with glucose as the sole carbon source as the defined environment for the first case of this study. Then, by compiling experimental and genome-scale model generated essential gene sets, we generated a list of essential genes for growth in this specific environment (Figure 1A, Supp. Table 1, see methods). Once the case specific gene essentiality is defined, we analysed the TF-gene interactions compiled in RegulonDB ^15^. After determining gene essentiality and TF-gene regulatory interactions, we analyse the sub-network of interactions of each TF (Figure 1B) looking for dispensable TFs, defined as those that do not activate the expression of any essential gene. According to our analysis, 156 from the 200 TFs contained in the regulatory network can be eliminated (Figure 1C). Since our goal is to reduce the hedging proteome, out of the 156 dispensable TFs we select as candidates for non-essential function reduction those 34 TF’s with at least one unique (meaning it is not activated by any other TF) positive regulated gene (Supp. Table 2) (See methods); this gives the certainty of silencing at least one gene.

### Integration of proteomic data and the TF-Gene regulatory network

We determine the proteome associated to each non-essential TF in our network integrating a quantitative proteomic data set^17^, that provides protein copy number per cell under 22 different growth conditions with 95% of proteome coverage (by mass). Here we define two emerging properties derived from the quantitative proteomics data integration: the Proteomic Load of a gene (P_L_) in fg of protein per cell (Figure 2A) and the Proteomic Balance (PB) of a TF resulting from the summation of the P_L_ of the genes that would result silenced or activated by the elimination of a TF (Figure 2B). P_B_ is conceptually important to rank the TFs according to the size of the proteome they control, since it takes into account the net addition of protein mass (in fg of protein/cell) liberated when removing a TF. A graphic representation of the 34 candidate TF subnetwork illustrating the P_L_ all the possible targets to affect is shown in Figure 4A.

**Figure 2.**
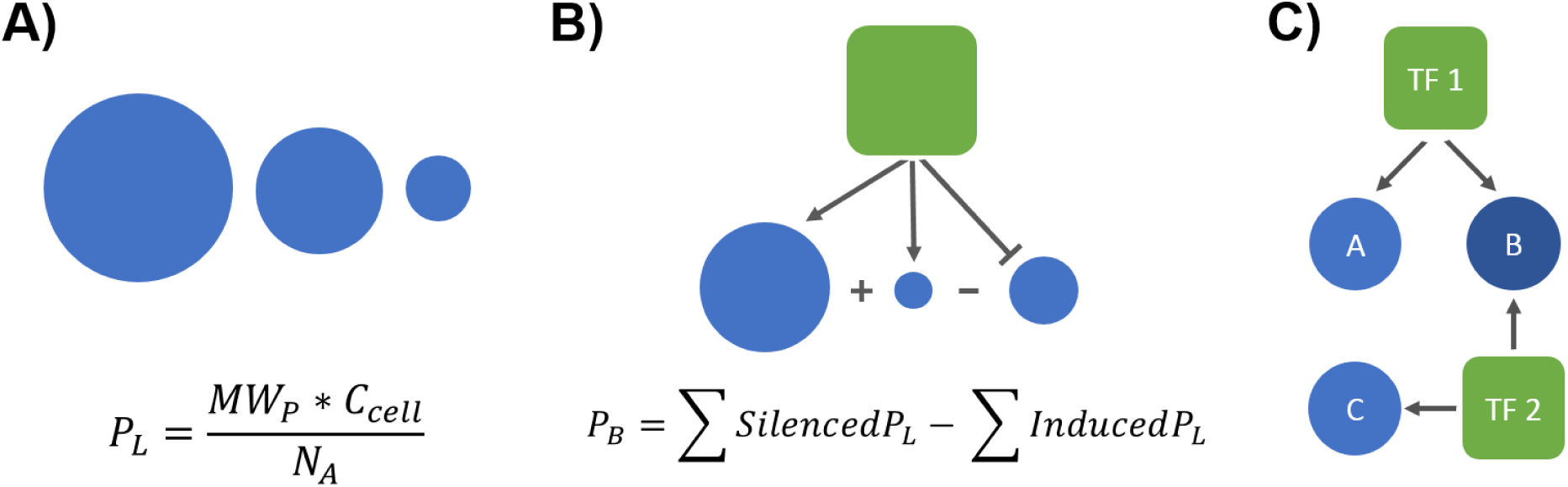
Emerging properties from proteomics data integration. **A)** Proteomic load of a gene (P_L_), this property is defined as the molecular weight of the protein (MW_p_) multiplied by the number of copies per cell (C_cell_) divided by Avogadro’s number (NA) (6.022×10^8^ fg equivalent), the more expressed the gene is, the more proteomic load it generates. **B)** Proteomic Balance of a TF (P_B_) which is defined as the sum of the P_L_ of the silenced genes minus the sum of the P_L_ of the induced genes. **C)** Schematic of a simple case of shared regulation in which removing both TFs silences all genes but this is not the case when the TFs are silenced individually.

**Figure 3.**
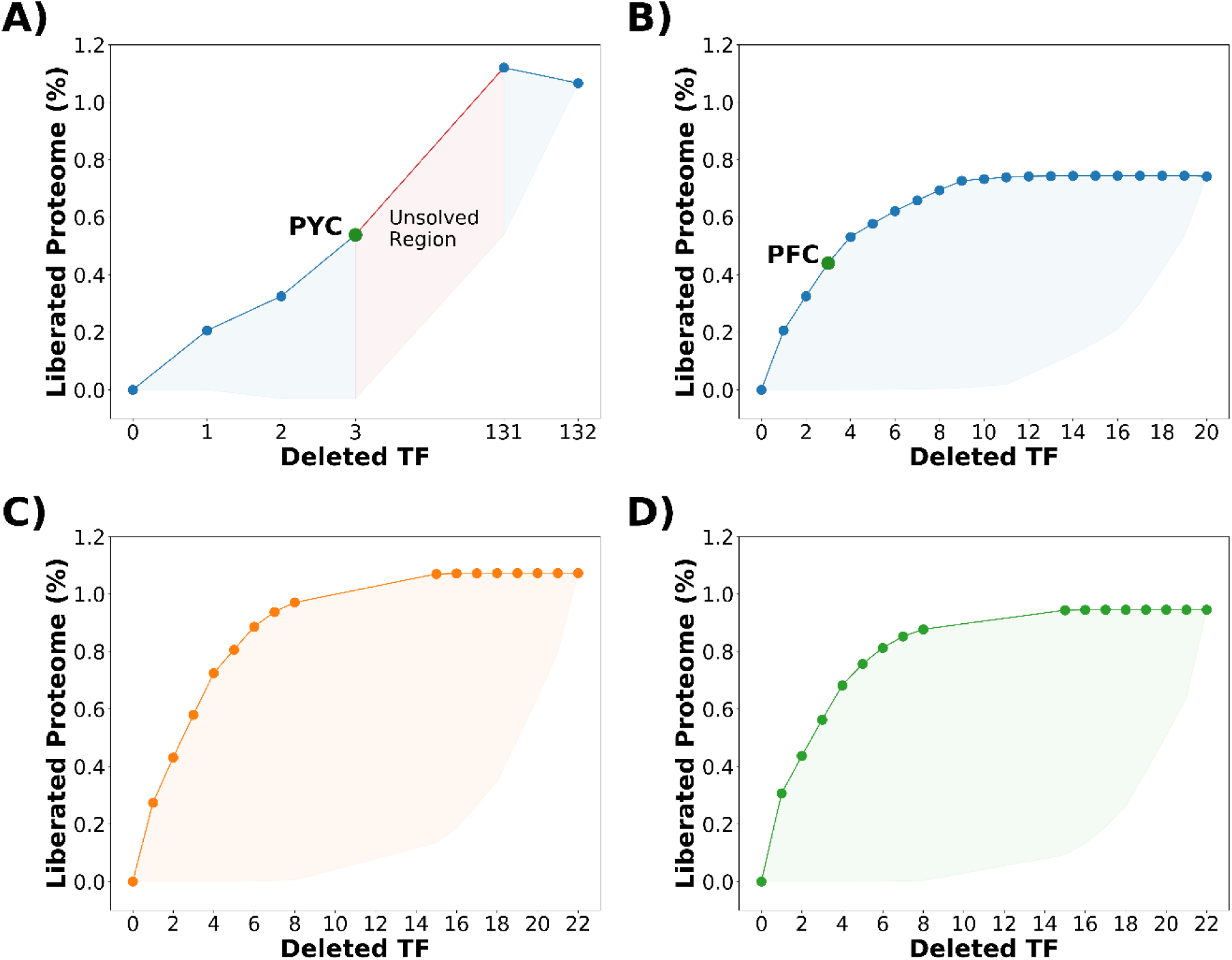
Proteome liberation calculations using ReProMin. **A)** Potential optimization landscape corresponding to the ST case; the solved region is shown in blue while the unsolved region in red, PYC mutant location in the landscape is shown with a green circle. Solved optimization landscape for UT case in **B)** glucose, PFC mutant location in the landscape is shown with a green circle, **C)** galactose and **D)** acetate.

**Figure 4.**
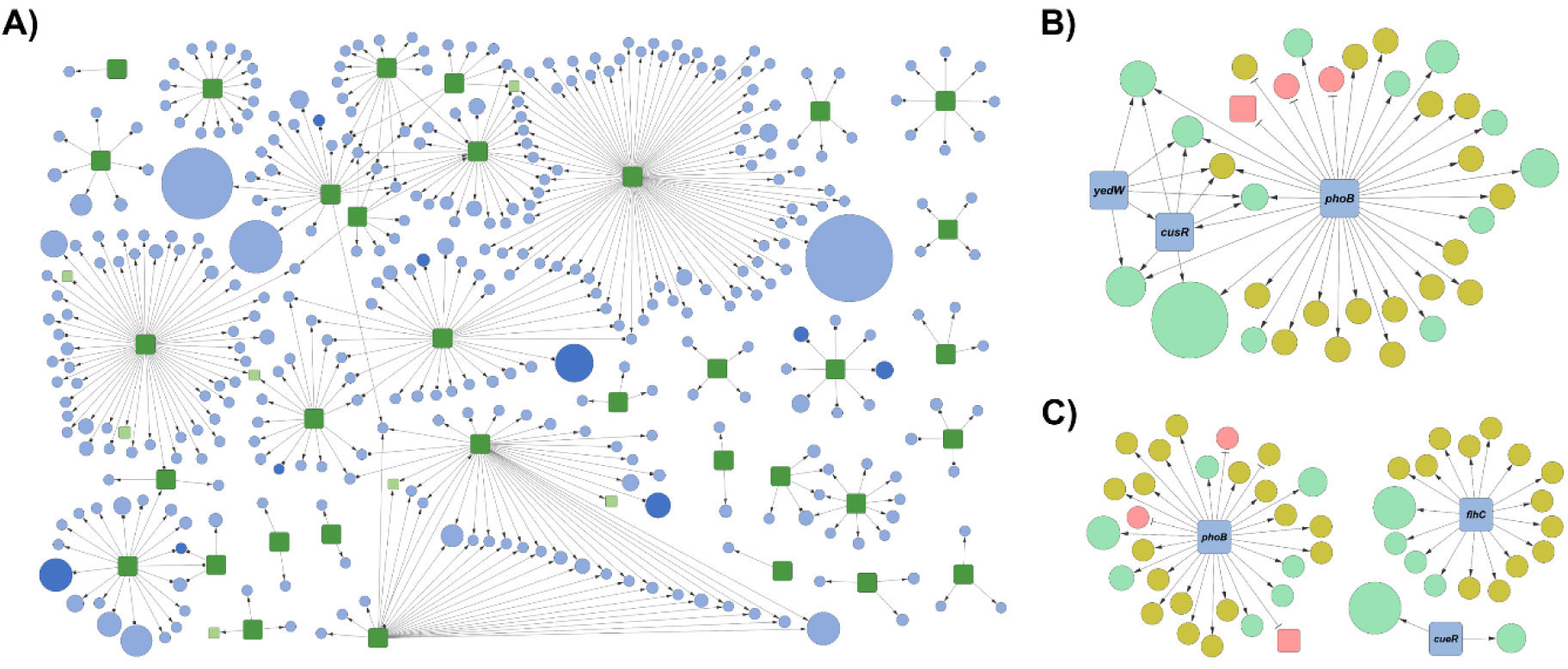
TF-Gene network and PL representation. **A)** Subnetwork of interactions corresponding to the 34 candidate TF representing all the potentially affected targets; green squares represent candidate TF, while light blue circles represent dispensable genes, dark blue circles are essential genes and light green squares are dispensable TF, circle size is proportional to the PL of the gene. Subnetwork of predicted regulated targets of the PYC **(B)** and PFC **(C)** mutants. In both cases, green circles represent predicted silenced targets, red circles predicted induced targets and yellow circles genes with no proteomic coverage; the size of the circles is proportional to the PL of the target. High resolution versions of the sub networks are available in the supplementary material (Supp. figs. 3 - 5).

### Computational search of minimal TF eliminations for the release of the maximal hedging proteome

Even though *E. coli* is one of the most studied organisms and its TRN has been widely investigated, only half of its genes have regulatory information (RegulonDB). In order to prevent detrimental effects on gene expression due to our incomplete knowledge of the regulatory network, we searched the smallest combination of TFs that liberate the greatest amount of resources. We observed that many TFs have shared target genes (Figure 2C), in fact, many of them are part of a simplified version of a Dense Overlapping Regulon (DOR) network motif^18^, meaning that a particular combination of TFs is needed in order to “fully” silence these targets. Due to the size of the landscape of potential phenotypes resulting from the combinatorial TFs deletions, we developed a computational tool to assist with the design of mutant strains. We called our tool ReProMin (**Re**gulation based **Pro**teome **Min**imization). ReProMin tool uses the previously described TF-Gene interaction network; quantitative proteomic data and a list of candidate non-essential TF to find the n-combination of mutations that silences the higher proteomic load (see methods).

In order to test our method (ReProMin), we first performed calculations using data from the glucose minimal media condition on which we defined gene and TF essentiality. Calculations were made considering two cases, the Shared Target (ST) case: considering all non-essential TFs with positive P_B_ (132 TF), which takes into account some TFs with no unique regulated genes. And the Unique Target (UT) case: considering candidate TFs with positive P_B_ (20 TF). Our computational tool was able to solve up to triple TF combinations for ST case and up to 20 TF combinations for UT case (Figure 3A and B).

For the ST case, calculations revealed that the elimination of all the non-essential TFs theoretically would liberate up to 1.06% of total proteome. However, up to 0.53% of total proteome can be released by removing a top combination of three TFs. For the UT case, the elimination of all 20 candidate TFs would liberate up to 0.72% of the proteome, and our simulations show that there is not a significant improvement in resource release after the elimination of 8 TFs.

We tested the accuracy of ReProMin predictions in other conditions for which proteomic data is available, such as growth on galactose and acetate minimal media. In this case we used the rich media essentiality gene set (see methods) and for the environment specific genes we performed essentiality simulations with a genome scale metabolic model in the corresponding growth condition (see methods). As a result, we obtained 164 and 166 non-essential TFs for galactose and acetate respectively and found that most of the identified TFs are shared among the three evaluated (glucose, galactose, acetate minimal media) meaning that they are all non-essential for minimal media growth with those carbon sources (Supp. fig. 2). In both cases we identified 23 candidate TFs that belong to the UT case and have a positive P_B_. Proteome liberation calculations were made using these subsets of candidate TFs. Our predictions show that we can release 0.88% and 0.81% of the total proteome in galactose and acetate, respectively, with the deletion of all these TFs (Figure 3C and D).

### Generation of combinatorial strains

Based on our ReProMin predictions, two triple knockout strains were generated, for the ST case: PYC (Δ*phoB* - phosphate scavenging system, Δ*yedW* - unknown gene, Δ*cusR* - copper efflux system) with a P_B_ of 1.3 fg representing 0.53% of the total proteome in glucose (Figure 3A), is a particular case of shared regulation where most of the target genes are only silenced by the deletion of all the three TFs together (a graphical representation of its TF-gene network is presented in Figure 4B). For the UT case: PFC (Δ*phoB* - phosphate scavenging system, Δ*flhC* - flagella master regulator, Δ*cueR-* copper efflux system) with a P_B_ of 1.08 fg representing 0.44% of the total proteome in glucose (Figure 3B), has a higher grade of confidence in the design than PYC due to simpler regulatory subnetwork (Figure 4C). We also generated a strain, using an intuitive approach, in which we eliminated three TFs that regulate non-growth related functions. The resulting strain is called FOG (Δ*fliA* - flagella sigma factor, Δ*oxyR* - oxidative stress master regulator, Δ*gadE* - acid resistance regulator). The FOG strain was not generated by our design pipeline; therefore the regulatory interventions may affect some important functions and it was used as a control to compare to our designed strains.

### RNA-seq analysis confirms the high specificity of introduced mutations

The predictive power of ReProMin depends on the accuracy of the interactions compiled in the *E. coli* TRN. We validated the predictions for the PFC mutant strain by comparing its transcriptome profile obtained by RNA-seq to that of the wild-type (WT). This experiment aims at determining the degree of success in gene silencing at the transcriptional level, and at assessing other possible transcriptional perturbations resulting from our regulatory modifications. Results show that no transcripts corresponding to the three deleted TFs were detected in PFC (Figure 5A). By mapping the fold change obtained in the analysis to the predictions of the computational tool, it is possible to visualize the impact at the transcriptional level of the missing regulators on their targets (Figure 5C). Four targets associated to *flhC*, corresponding to genes forming the flagella (*flgB, flgC, flgE* and *flgG*) were completely silenced; furthermore, all the other flagella-related genes also registered a decrease on their expression. Regarding *phoB*, two targets were successfully silenced (*phnI* and *phnL*), both genes belong to an operon that is induced under phosphate starvation and is required for use of phosphonate and phosphite as phosphorous sources ^19^, many other targets related to this operon also reduced their expression. On the contrary, *phnK* present in the same operon was surprisingly overexpressed. We were unable to map any transcripts to six genes belonging to the previously mentioned operon, which may not be entirely expressed in the absence of phosphate starvation. Furthermore, *phoR* (part of the *phoB-phoR* two-component system) also reduced its expression. Finally, both targets of *cueR (copA* and *cueO*) also decreased drastically their abundance.

**Figure 5.**
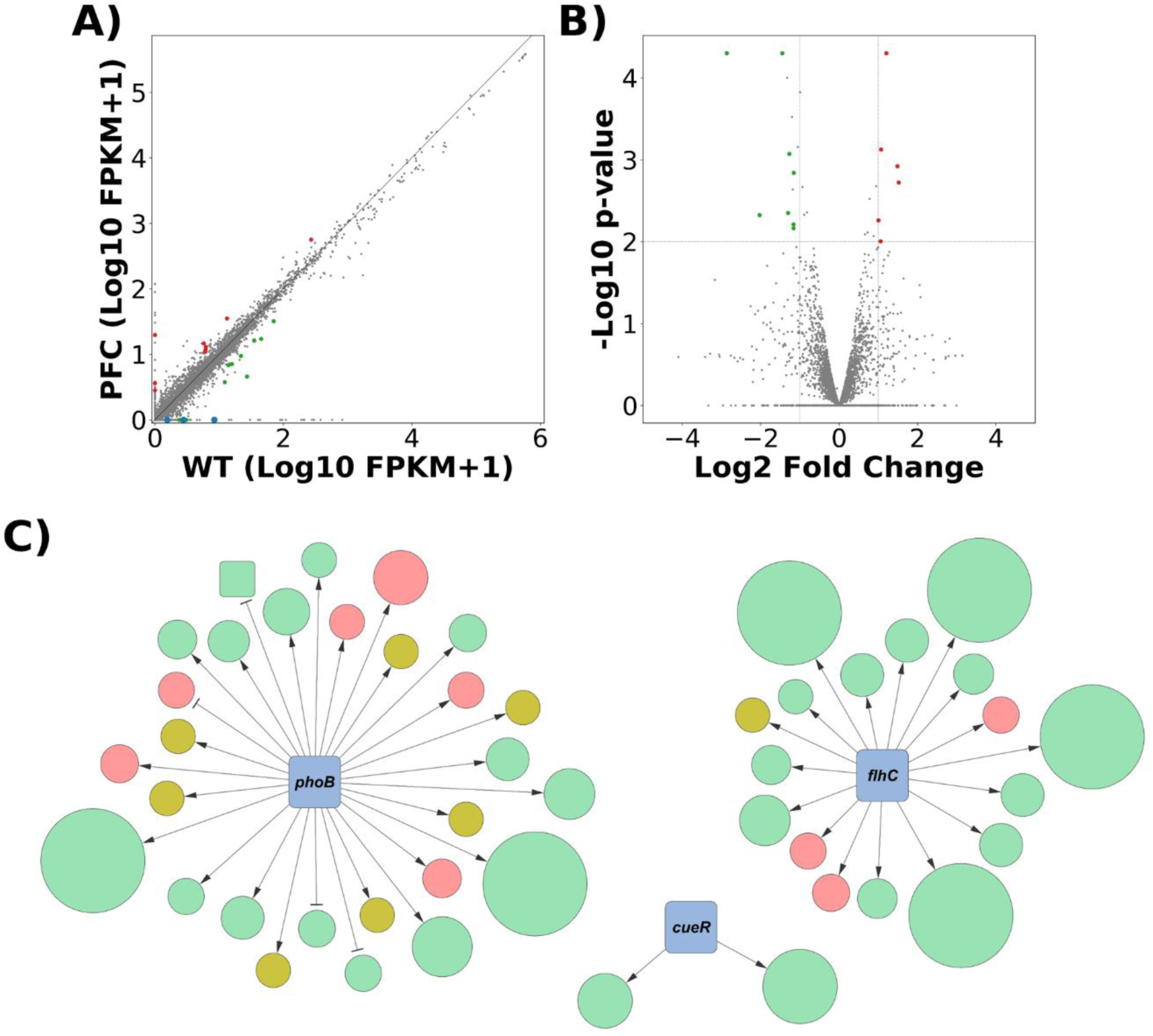
Transcriptomic analysis of the designed strain. **A)** Correlation plot for PFC and WT strains transcripts. Blue dots represent the three deleted TFs. **B)** Volcano plot showing differential gene expression. In both cases, statistically significant genes are highlighted. **C)** Integration of transcriptomics with computational tool predictions. The size of the circle corresponds to the fold change of each target (the largest circles represent fully silenced genes), in all cases green circles represent targets releasing resources (down regulated), red circles represent targets generating burden (up regulated) and yellow circles targets that were not expressed. High resolution version of the sub network is available in supplementary material (Supp. fig. 6).

Regarding the accuracy of our ReProMin predictions, 28 genes of 47 predicted silenced genes were silenced at different levels, whereas 11 predicted silenced genes presented higher expression values than the WT strain. Finally, transcripts of 8 predicted targets were not found in either strain. These observations show that in 72% (28 of 39 measured genes) of the cases the predictions of the computational tool were accurate (Supp. fig. 7).

Additionally to the designed transcriptional changes, we found 17 genes differentially expressed (8 down regulated genes and 9 up regulated (log2 Fold Change ≥ 1 or ≤ −1 and p-value ≤ 0.05) (Figure 5B) (Supp. Table 3). This RNA-seq analysis shows that besides the intended transcriptional changes, few off-target effects were identified at the transcriptomic level in the PFC strain.

### Phenotypic evaluation revealed reduced burden, increased cell yield and production budget for designed strains

Our three ReProMin generated mutants (ST case, UT case, and control) were evaluated in rich (LB) as well as in minimal media containing three different carbon sources (acetate, galactose, glucose). The computationally designed strains (PYC and PFC) showed neither growth defects nor increase in the growth rate or biomass yield in any of the four conditions tested. On the contrary, the control strain (FOG) showed growth defects in all growth condition tested (Supp. fig. 8). For glucose minimal media, we also evaluated the effect of recombinant protein production using a plasmid expressing a genetic circuit with two fluorescent reporters (Figure 6A) ^20^. The burden caused by carrying a plasmid is reflected as a decrease in the growth rate in all tested strains (Supp. fig. 9A); this decrease is higher when the plasmid is expressing the genetic circuit, however the burden displayed by both ReProMin designed strains is lower compared to the wild-type counterpart. Additionally, the PFC strain also showed a higher final biomass production (Supp. fig. 9B). It has been described that the expression levels of two protein reporters encoded on the same plasmid but without a regulatory connection between them is captured by a linear relationship which can be interpreted as an isocost line, a concept used in microeconomics to describe how two products can be bought with a limited budget, so the more is used on one, the less can be used on the other. These lines represent the boundary of the production budget of a given strain and condition (Figure 6B) ^20^. We obtained the isocost lines at balanced growth (∼5 h.) determined by two different methods: mean plate reader fluorescence and mean fluorescence measured by flow cytometry. The line corresponding to PFC strain show a parallel upward shift compared to the WT strain, which represents an increase of 9% in absolute fluorescence (p < 0.01) (Figure 6C) and 12% in mean fluorescence per cell (p < 0.01) (Supp. fig. 10), this difference is increased at the stationary phase of the culture (∼24 h.) were higher maximal biomass is achieved and the quantity of recombinant protein is increased up to 18% (Figure 6D).

**Figure 6.**
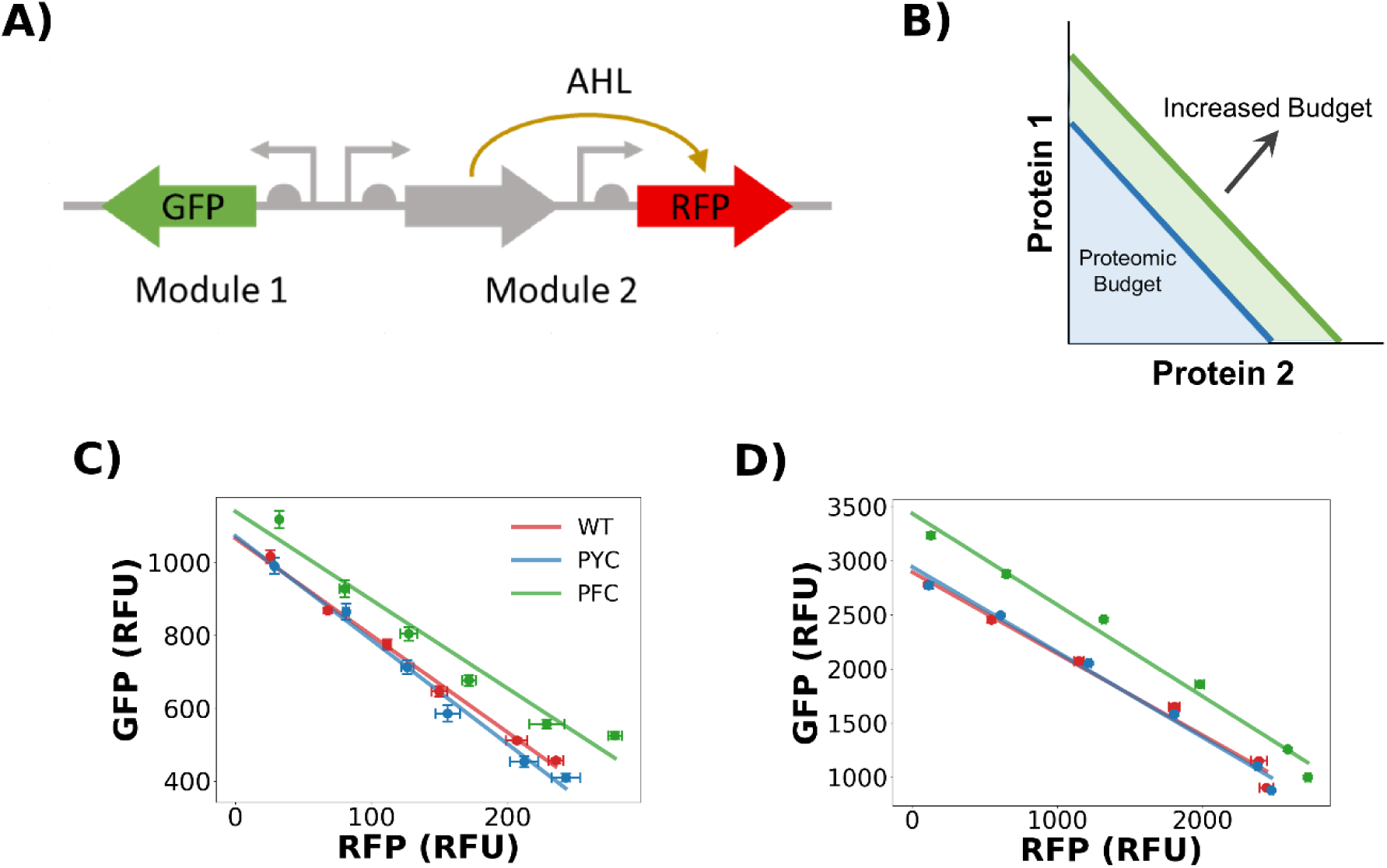
Synthetic circuit characterization. **A)** Schematic of the gene circuit evaluated, which encodes the fluorescent reporters GFP and RFP: the first is constitutively expressed, while the latter is under the control of an N-acyl-homoserine lactone (AHL) inducible promoter. **B)** When plotting the expression of one protein against the other at different levels of induction, an isocost line is obtained. The size of the area below the line represents the total proteome budget dedicated to the circuit, a parallel upward shift in the line represent an increase in the budget. Isocost lines of the designed strains showing absolute fluorescence during **C)** balanced growth and **D)** stationary phase. Each point represent the red reporter (x axis) plotted against the green reporter (y axis) in an increasing inductor concentration (1.25, 2.5, 5, 10, 20 nM AHL) and represents the mean of three replicates across three different experiments. A linear regression was used to fit the points to a line.

**Figure 7.**
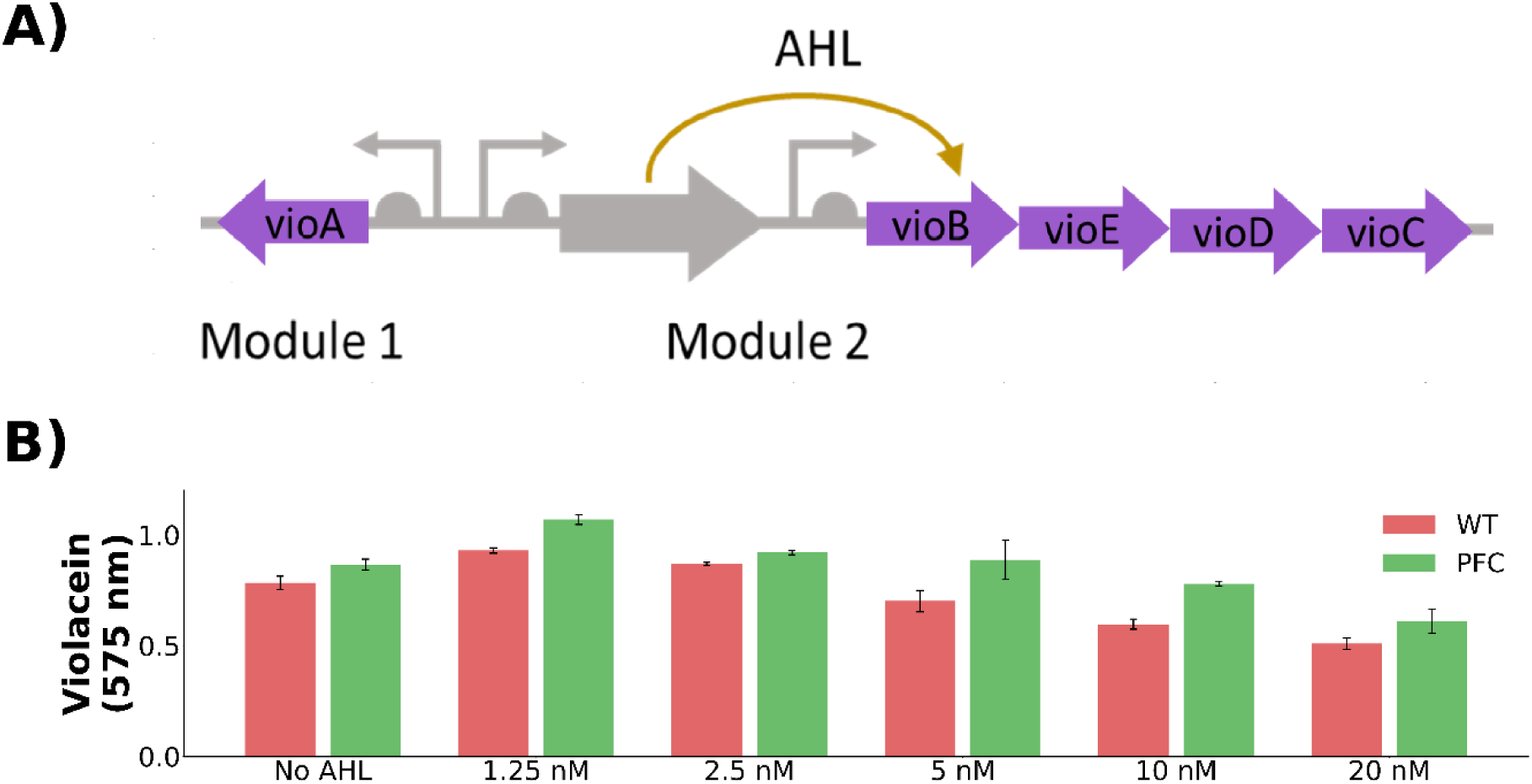
Violacein Pathway Evaluation. **A)** Schematic of the pathway of violacein biosynthesis. **B)** Total violacein production using 2 g/L tryptophan after 24 h in the presence of increasing inducer concentrations; each value represents the mean of three replicates across three different experiments.

### Expression of an engineered metabolic pathway: violacein production

We tested the ability of our engineered strain for synthesizing the molecule violacein as a proof-of-concept for applications of our method in metabolic engineering. Violacein is a pigment from *Chromobacterium violaceum* endowed with many biological activities (antibacterial, antiviral, anti-parasite) and has recently gained importance in the industrial field especially for applications in cosmetics, medicines and fabrics ^21^. Violacein is synthesized in a five-step metabolic pathway using tryptophan as a precursor. Here we used the violacein pathway plasmid reported by Darlington *et al*., 2018, where the five genes for violacein biosynthesis are arranged in two operons, one consisting of *vioA* constitutively expressed, while the rest of the pathway encoded by the *vioBCDE* genes is under the control of an AHL inducible promoter ^13^ (Figure 7A). This construction follows the same principle as the previous circuit so the more of one module is produced, the less of the other is expressed due to the competition for limited resources for gene expression. However, in this case the number of genes in each module is different and code for actively metabolic enzymes with different kinetic properties, which results in differential violacein biosynthesis.

We evaluated violacein production after 24 hours in the wild-type and PFC strains using M9 glucose medium supplemented with tryptophan (2.0 g/L) and AHL (1.25, 2.5, 5, 10, 20 nM) for induction. PFC showed a mean increase in violacein production of 18% (Figure 7B), additionally we found that the maximum production is achieved with just a minimum quantity of inducer (1.25 nM) indicating that is crucial to have a balanced expression of the pathway with the right amount of each module to maximize the synthesis of the final product. Similarly to our observations on fluorescent protein production, the increase in violacein production shows that our approach can be harnessed to increase the production of metabolites from costly heterologous metabolic pathways.

## Discussion

Gene regulatory networks are robust and can be severely rewired with interesting phenotypic outcomes^22^, thus they are a perfect rational engineering target for synthetic biology applications. In this work we prove that the definition of an essential gene set together with regulatory network information allows the identification of TFs whose elimination leads directly to silencing proteome fractions that are not used in a particular condition. We show that by eliminating hedging proteome activators we can release resources and increase cellular capacity for engineered functions. In agreement with the presented ME-model simulations reducing the unused protein fraction, our designed strain shows higher proteomic budget, measured by the isocost lines, and a higher capacity to produce a metabolite from a heterologous pathway. Furthermore, by comparing our strains with an intuitive control strain, we show that inaccurate TF elimination results in detrimental effects on growth, maximal biomass and protein production (Supp. figs. 9 and 11). These findings indicate that the elimination of a combination of TFs is not a trivial process; it may affect essential functions and introduce phenotypic defects. Our method shows good accuracy in terms of the obtained gene expression changes measured by RNAseq, despite our limited knowledge of the regulatory networks. In addition, the regulatory data available is condition dependent, what limits the predictive power of our method, since we need to assume that regulatory interactions are present at all times. We anticipate that developments in high throughput technologies (such as Chip-seq) combined with novel computational approaches^23–25^ will enable the fast generation of complete regulatory networks and the application of our method to even non-model organisms. Several approaches have been applied for resource allocation optimization in bacterial host engineering. Genome minimization has been mainly done by large scale genetic interventions whose outcomes are difficult to predict and do not show greater genome stability ^26,27^. Adaptive Laboratory Evolution (ALE) has showed great success, especially to identify functions not related to growth ^28^, however, it selects for fast growing strains which not necessarily result in the best production phenotypes. Moreover, the underlying selection mechanisms in ALE are normally not known therefore its effects are not predictable ^29^. Genome scale models, such as the ME-model, may also be used to find the proteomic cost and fitness benefit of gene expression, thus to aid in the design of proteome allocation, however kinetic data of each protein is needed ^30^ and its scope focuses on growth related functions. There are only a few reports describing regulatory approaches to improve production phenotypes, such as the global Transcriptional Machinery Engineering (gTME) ^31^, but none of them followed a rational approach. The methodology presented in this work is a novel strategy for proteome optimization with minimal genetic interventions which overcomes the serious limitations of deleting large regions of the genome; it is a flexible pipeline which can be applied to other growth and production conditions and also to different organisms where sufficient information is available. This work shows the potential of rational design of biological systems over the predominantly used trial and error approaches.

## Materials and Methods

### 1) ME-Model Simulations

All simulations were done using model iJL1678b ME ^14^. The corresponding transcription and translation reactions for recombinant protein (GFP) production were manually added to the model using standard methods. Unused protein fraction and flux through the recombinant protein production are changeable variables in the ME-Model that affect predicted growth rate and proteome composition, the values of these two variables were systematically changed in the ME-Model to assess their effect on growth rate (UPF = 0.36, 0.30, 0.25; Flux = 0, 0.001, 0.002, 0.0025, 0.0030, 0.0035, 0.0040), all other model parameters were set as default. Proteome sectors were classified according to O’Brien *et al*., 2016.

### 2) Definition of the essential gene list

To compile the essential gene list in the glucose minimal media condition we combined five different datasets from different sources. Three of them were experimentally generated using different methods of gene disruption: A) random transposon mutagenesis using M9 with glucose as growth condition (Tn-seq)^32^, B) removing large fragments of the chromosome using a homologous recombination system in rich medium (LB) ^33^ and C) the updated list of the mutants of the Keio collection that are lethal, the collection was generated using rich medium ^34,35^. Two gene lists were generated *in silico* using simulations of genome scale metabolic and expression models capable of predicting gene expression needs in a particular condition: D) genes that are essential for growth in M9 with glucose using iOL1554-ME model ^36^ and E) genes that are essential for growth in the metabolic model iJO1366 and also experimentally in M9 with glucose ^37^. Within the compiled list, genes exclusively belonging to the Tn-seq and the glucose minimal media ME-model simulations gene lists were considered conditionally essential, as these gene lists were originally generated using M9 with glucose as the growth condition, while the rest of the genes were classified as core essential for our purposes.

For the cases of galactose and acetate minimal media conditions we performed gene essentially analysis with the iML1515 model ^38^ in COBRApy ^39^.

### 3) Identification of candidate regulators and combinatorial analysis

We sorted the TF/gene interactions from RegulonDB (version 8, regulondb.ccg.unam.mx), discarding all the sigma factors-gene interactions. Next, we classified as essential all TFs that activate at least one essential gene (from our M9 glucose condition gene list) and as non-essential all TF’s that do not activate any essential genes. Then we analysed the sub-network of interactions of each non-essential TF by numerically analysing the output level of each TF (TF_OUT_), which is classified into positive and negative output (TF_OUT+_ and TF_OUT-_) representing positive and negative regulated genes respectively and the degree of entry of each regulated gene (GENE_IN_) in turn also divided into positive and negative (GENE_IN+_ and GENE_IN-_). We defined as candidate for proteome reduction all those TFs that activate at least one unique gene, which numerically meet the following condition and represent a simplified version of a SIM:

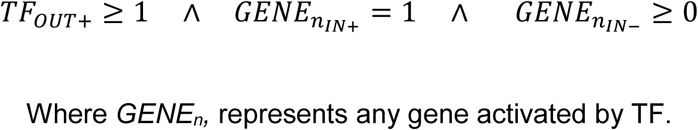

On the other hand, the proteomic dataset previously described ^17^ was used to calculate the Proteomic Load (P_L_) of each gene and Proteomic Balance (P_B_) of each TF according to the equations in Figure 2.

The combinatorial analysis was achieved as follows, given a list of TFs, we created and tested all the possible N combinations. Next, the total number of silenced and induced genes for each combination was determined following the next criteria: for every gene involved in the combination, we subtracted one from the value of GENE_IN+_ for each TF that regulates the target gene positively and one to the value of GENE_IN-_ for each TF that regulates the target gene negatively. At the end of this process a gene was considered silenced if:

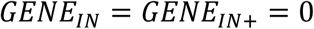

or induced if:

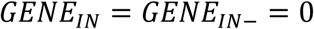

Finally, the P_B_ of each combination tested was calculated and ranked. The full computational set of tools coded in Python and datasets used in the analysis are available in the following repository (https://github.com/utrillalab/repromin). Cytoscape software version 3.7 ^40^ was used to plot the network representation of the data.

### 4) Generation of combinatorial knock-out strains

The combinatorial mutants were generated by sequential P1 phage transduction from the individual knock-out strains of the Keio collection according to the protocol described by Miller (1992) ^41^. The removal of the kanamycin resistance cassette before each transduction was done using the pE-FLP plasmid (Addgene plasmid #45978), pE-FLP was a gift from Drew Endy & Keith Shearwinand. Each knock-out strain was confirmed by PCR using primers flanking each gene. In all experiments *Escherichia coli* BW25113 was used as the WT background. The characteristics of the strains, plasmids and primers used in this study are described in the supplementary material (Supp. Table 4).

### 5) RNA sample extraction and sequencing

Strains were grown in 50 mL of M9 media with glucose (4 g/L) M9 media in 250 mL Erlenmeyer flasks cultures in an orbital incubator at 37°C (250 rpm). Cells were harvested in mid-log phase using the Qiagen’s RNAprotect bacteria reagent according to the manufacturer’s specifications. Cell pellets were incubated with lysozyme, SuperaseIn and protease K for 10 min at 37°C. Total RNA was isolated and purified using Zymo Research’s Quick-RNA kit according to the manufacturer’s specifications. All samples’ quality was inspected in a bioanalyzer RNA chip (Agilent). Starting with 10ug of total RNA of each sample, the removal of ribosomal RNA was done with the Ribominus kit by Invitrogen. For the construction of the libraries, the TruSeq Stranded mRNA Sample Prep Kit by Illumina was used, following the HT protocol. For sequencing a NextSeq 500 v2 was used, with a configuration of 2 × 75 paired-end read and 10 million reads per sample.

Reads were mapped to reference genome *E. coli* MG1655 (RefSeq: NC_000913.3) using aligner Bowtie2 (http://bowtie-bio.sourceforge.net/bowtie2). Final differential analysis was made using the Cufflinks library (http://cole-trapnell-lab.github.io/cufflinks). Genes with log2 Fold Change ≥ 1 were considered up regulated and ≤ −1 down regulated, considering a p-value ≤ 0.01.

### 6) Growth phenotype characterization

For the evaluation of growth in different carbon sources, the following conditions were used: glucose M9 medium (4 g/L), galactose M9 medium (3.2 g/L), acetate M9 medium (2.5 g/L) and LB. Cells were cultured overnight in the corresponding media. The next day the strains were diluted to an OD_600_ of 0.05 in fresh medium and 150 μl of the fresh culture were transferred to a transparent 96-well plate (Corning) and incubated at 37 °C with fast linear shaking in a microplate reader (Synergy 2.0, BioTek) for 24 hours, taking measurements for OD_600_ every 20 min. In all trials three replicates were included and the experiment was repeated independently on three different days.

The characterization of the growth kinetics was conducted using the algorithm developed by Swain *et al*., ^42^ with the default parameters. In all cases, when comparing the parameters obtained from the mutant strains against the wild type, the p-value was determined using a two-tailed Student t-test as the statistical significance reliability index.

### 7) Isocost circuit evaluation

Strains were inoculated into glucose M9 medium with gentamicin (20 ug/ml), and grown overnight. Next day strains were diluted to an OD600 of 0.05 in fresh glucose M9 medium containing *N*-acyl homoserine lactone (AHL, Sigma-Aldrich, St. Louis, MO, USA, final concentrations of 1.25, 2.5, 5, 10, 20 nM), then 150 μl of the fresh culture were transferred to a 96-well black plate with transparent bottom (Corning) and incubated as described above, taking measurements for OD_600_, GFP (ex., 485 nm, em., 528 nm) and RFP (ex., 590 nm, em., 645 nm). The characterization of the production kinetics of GFP and RFP was also done using the algorithm described above.

### 8) Flow cytometry measurements

For flow cytometry measurements, cell cultures were prepared as described above, but later grown in 24-well plates using 1 ml of medium. Every hour 50 μL aliquots were taken from each well and mixed with 150 μL of PBS, the volume of the wells was kept constant by adding fresh medium. Cell suspension was loaded into an Attune NxT Flow Cytometer (ThermoFisher, Waltham, MA, USA) and analysed for GFP (excitation 488 nm; emission 525/50 nm) and RFP (excitation 561 nm; emission 620/15 nm). For each sample 20,000 events were analysed and population means were estimated using the default software of the instrument.

### 9) Characterization of violacein-producing strains

The strains were inoculated into glucose M9 medium with gentamicin (20 ug/ml), and grown overnight. Next day strains were diluted to an OD_600_ of 0.05 in fresh glucose M9 medium containing AHL (1.25, 2.5, 5, 10, 20 nM) and tryptophan (0, 0.5, 1.0 and 2.0 g/L), then 150 μl of the fresh culture were transferred to a 96-well plate and incubated as described above. After 24 h the plate was centrifuged (13,000×g, 10 min), and the supernatant of each well was discarded. Violacein was extracted by suspending the pellet in each well in 200 µl absolute ethanol and incubating the plate at 95 °C for 10 min followed by pelleting cell debris (13,000×g, 10 min). Violacein present in the extract was determined spectrophotometrically at 575 nm in a microplate reader (Synergy 2.0, BioTek).

## Acknowledgements

Elisa Marquez-Zavala and Colton Lloyd for ME-Model simulations support. Yared Castillo-Franco for computational support and Georgina Hernandez-Chavez for technical support.

## Funding

UNAM–DGAPA-PAPIIT projects IA200716 and IA201518. Newton advanced Fellowship Project NA 160328. JJ, and JK acknowledge the support received from the Biology and Biotechnology Research Council (BBSRC) (grant BB/M009769/1) and European Union’s Horizon 2020 research and innovation programme for the project P4SB (grant agreement no. 633962).

## Author contribution

JU and GLP designed ReProMin; GLP developed computational methods and performed data analysis; GLP, JSMH carried out experiments; GLP, JK and JIJ analysed flow cytometry experiments, isocost lines and violacein production; JU supervised the study; JU, GLP and JIJ wrote the manuscript.

## Data availability

RNA-seq data from this study have been deposited in NCBI’s Gene Expression Omnibus (GSE 134335). The code to run ReProMin can be found at: https://github.com/utrillalab/repromin

## Conflict of interest

JU and GLP are inventors in a MX patent application filled by UNAM.

## Supplementary Figures

**Supplementary Figure 1.**
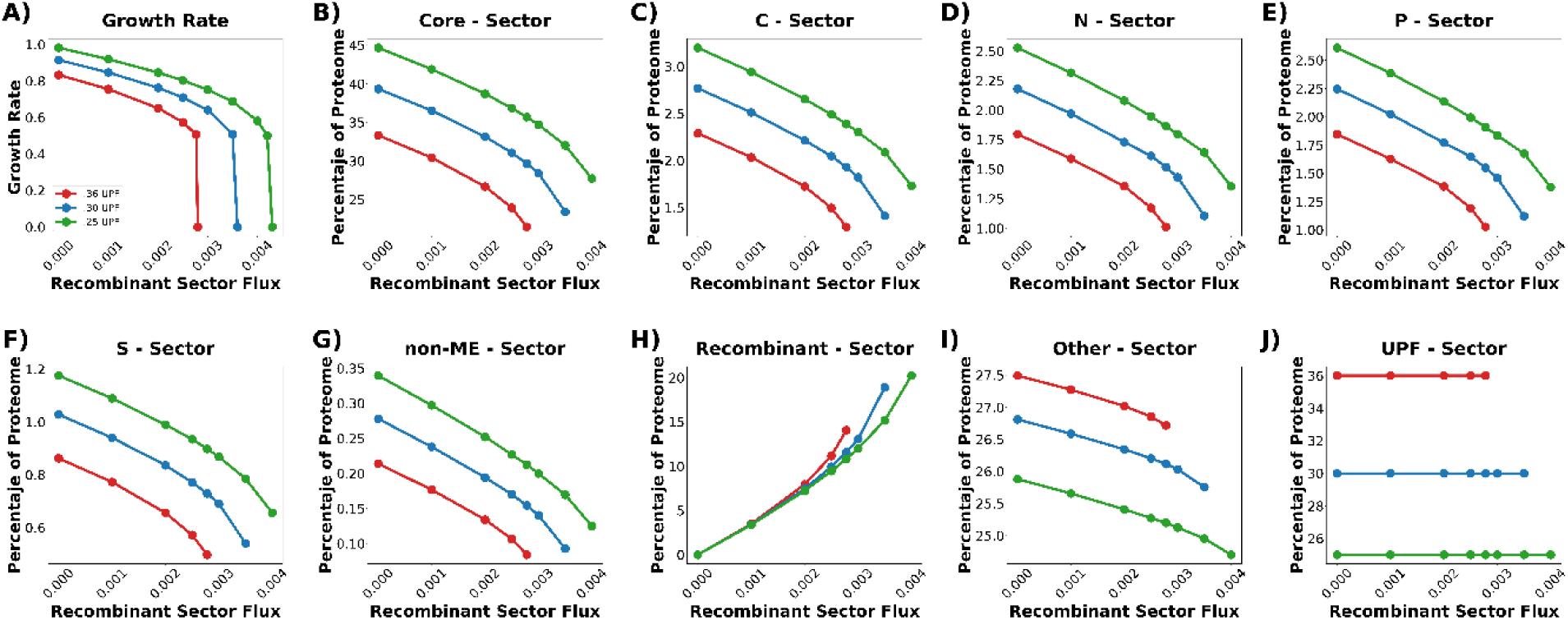
ME-Model simulations and proteome sector response. The expression of hedging functions has a proteomic cost that impacts growth and resource availability for the recombinant sector ^9, 10^. Genome scale models of metabolism and gene expression can be used to analyse the amount of resources devoted to those functions and predict engineering outcomes. Similar to the maintenance energy coefficient ^43^, the hedging proteome and other non-growth related (thus not modelled) functions are accounted for in ME-models as a part of the unused protein fraction (UPF). This proteome sector is comprised by functions not directly related to growth in the simulated environment; therefore those functions are not included in the model. The ME-model *i*JL1678b-ME ^14^ was used to simulate the effect of the reduction of the UPF and different expression levels of an unused recombinant model protein (GFP). The simulation shows an increased availability of cellular resources for recombinant protein production by reducing the UPF.

**Supplementary Figure 2.**
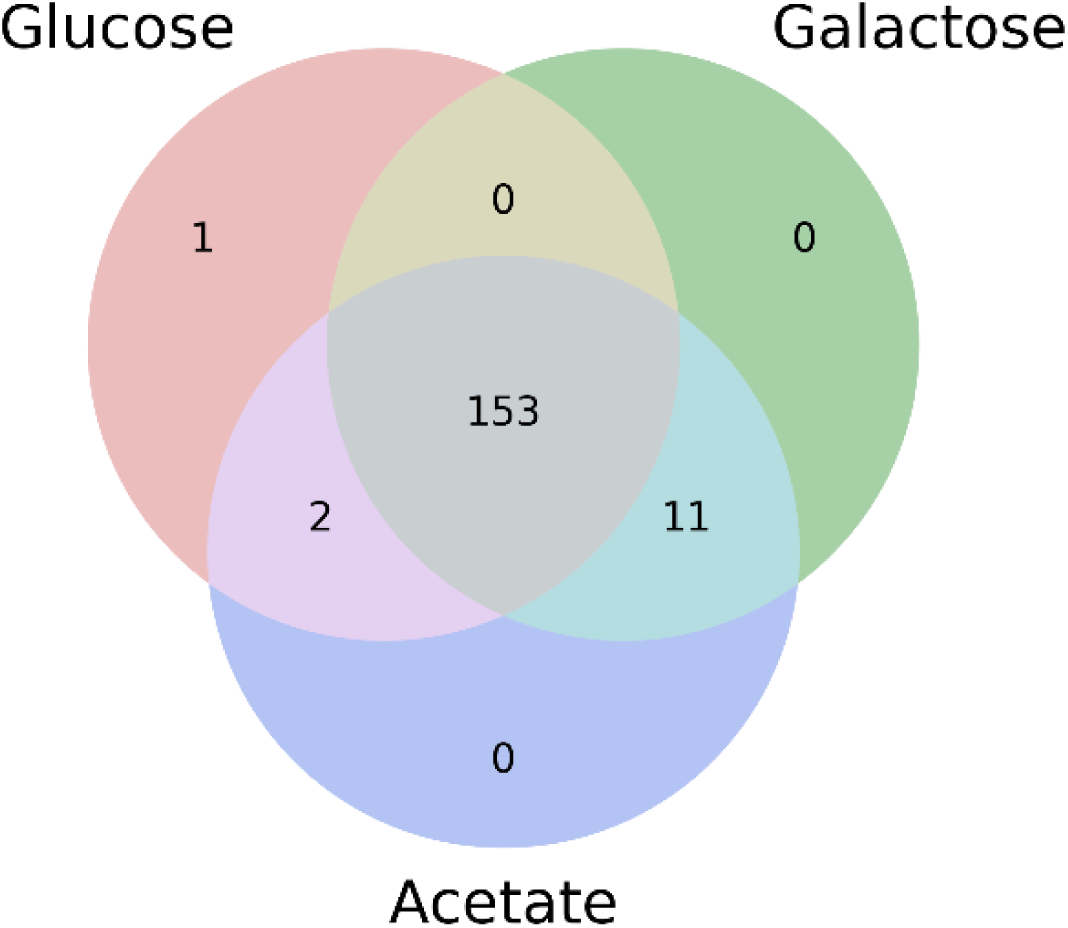
Non-essential TF distribution for the UT case across three growth conditions tested in this study.

**Supplementary Figure 3.**
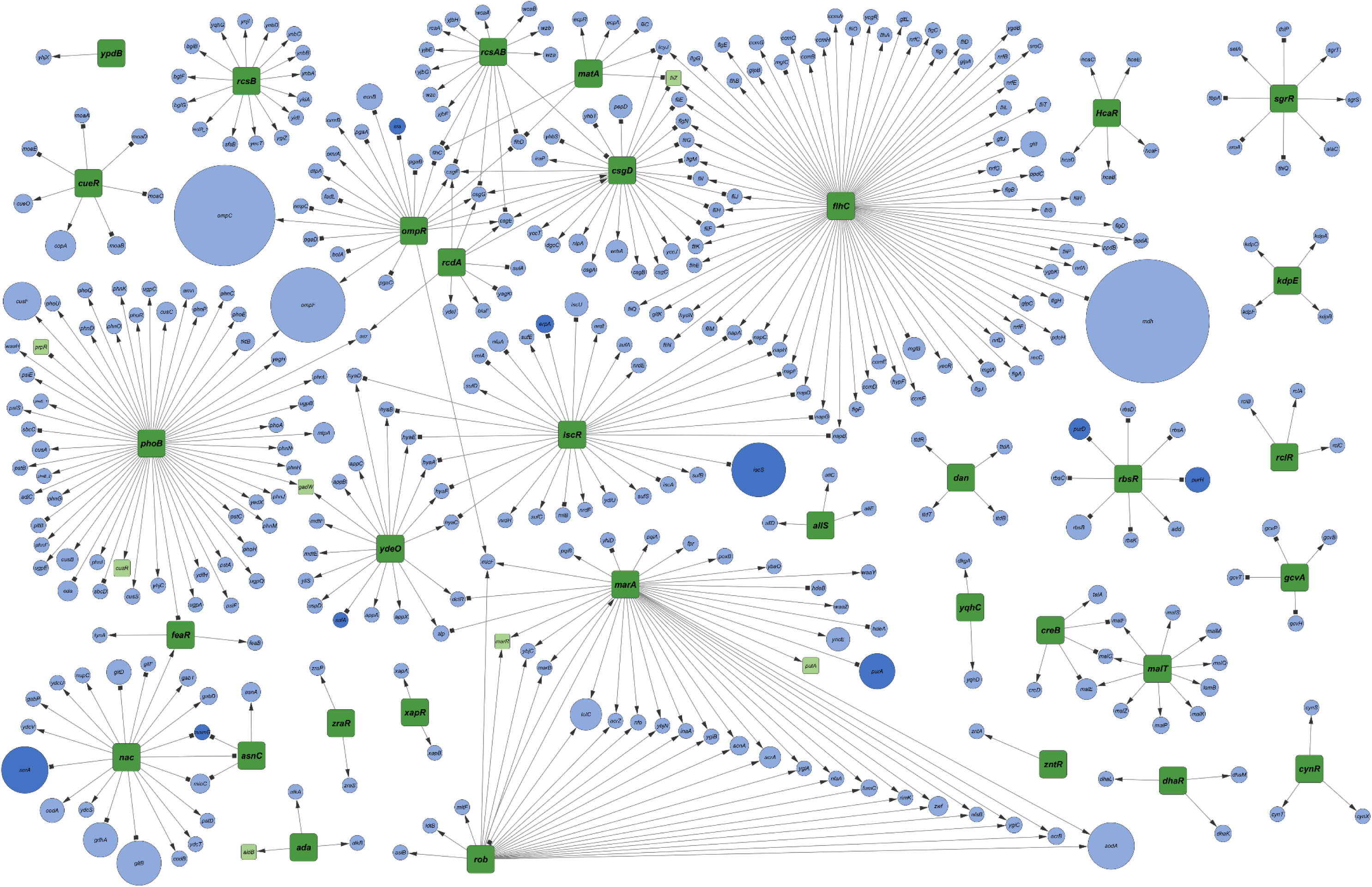
Subnetwork of interactions corresponding to the 34 candidate TF representing all the potentially affected targets; green squares represent candidate TF, while light blue circles represent dispensable genes, dark blue circles essential genes and light green squares dispensable TF, circle size is proportional to the PL of the gene.

**Supplementary Figure 4.**
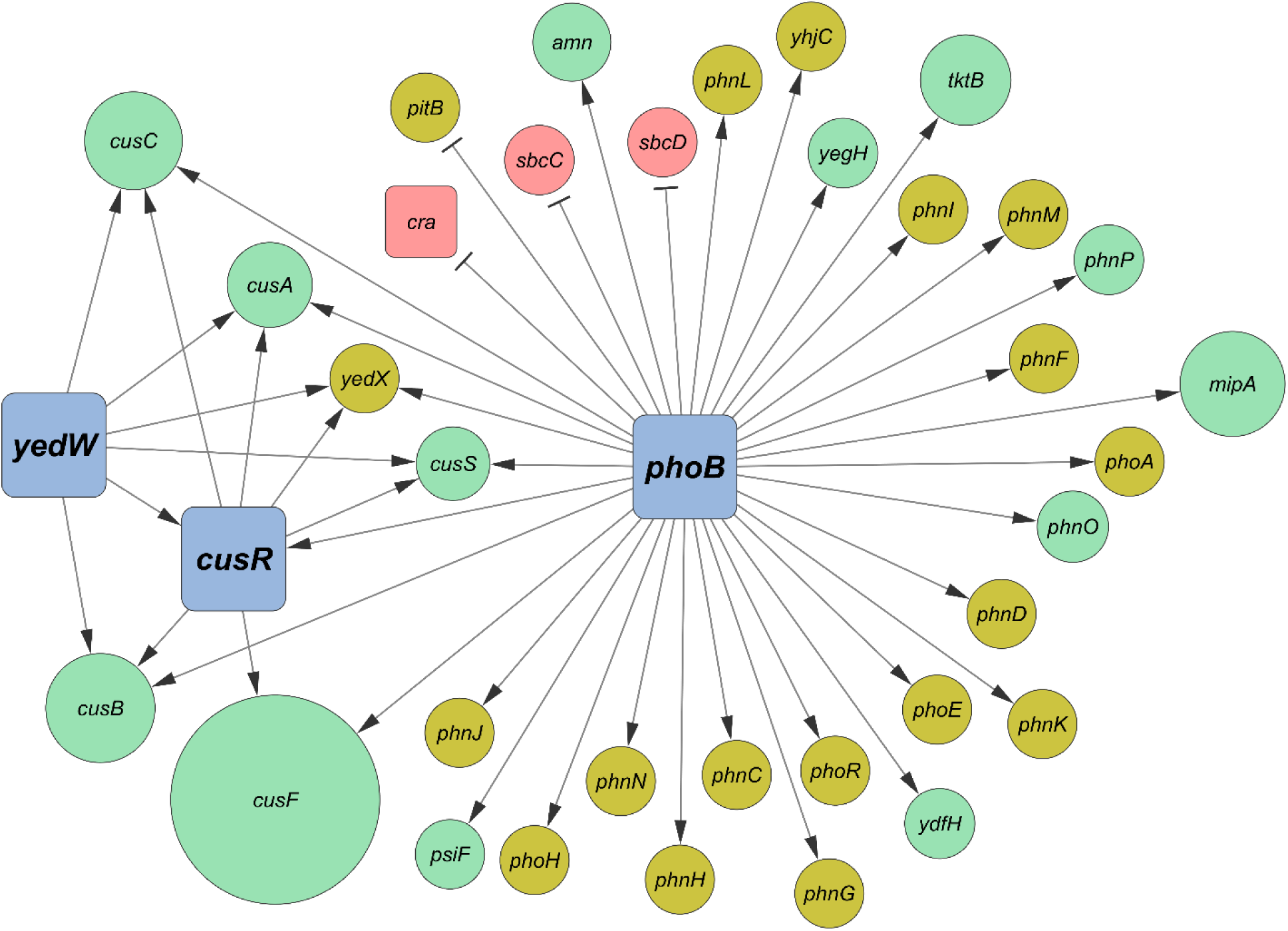
Regulatory subnetwork of predicted gene targets of the PYC mutant; green circles represent predicted silenced targets, red circles predicted induced targets and yellow circles genes with no proteomic coverage; size of the circles is proportional to the PL of the target.

**Supplementary Figure 5.**
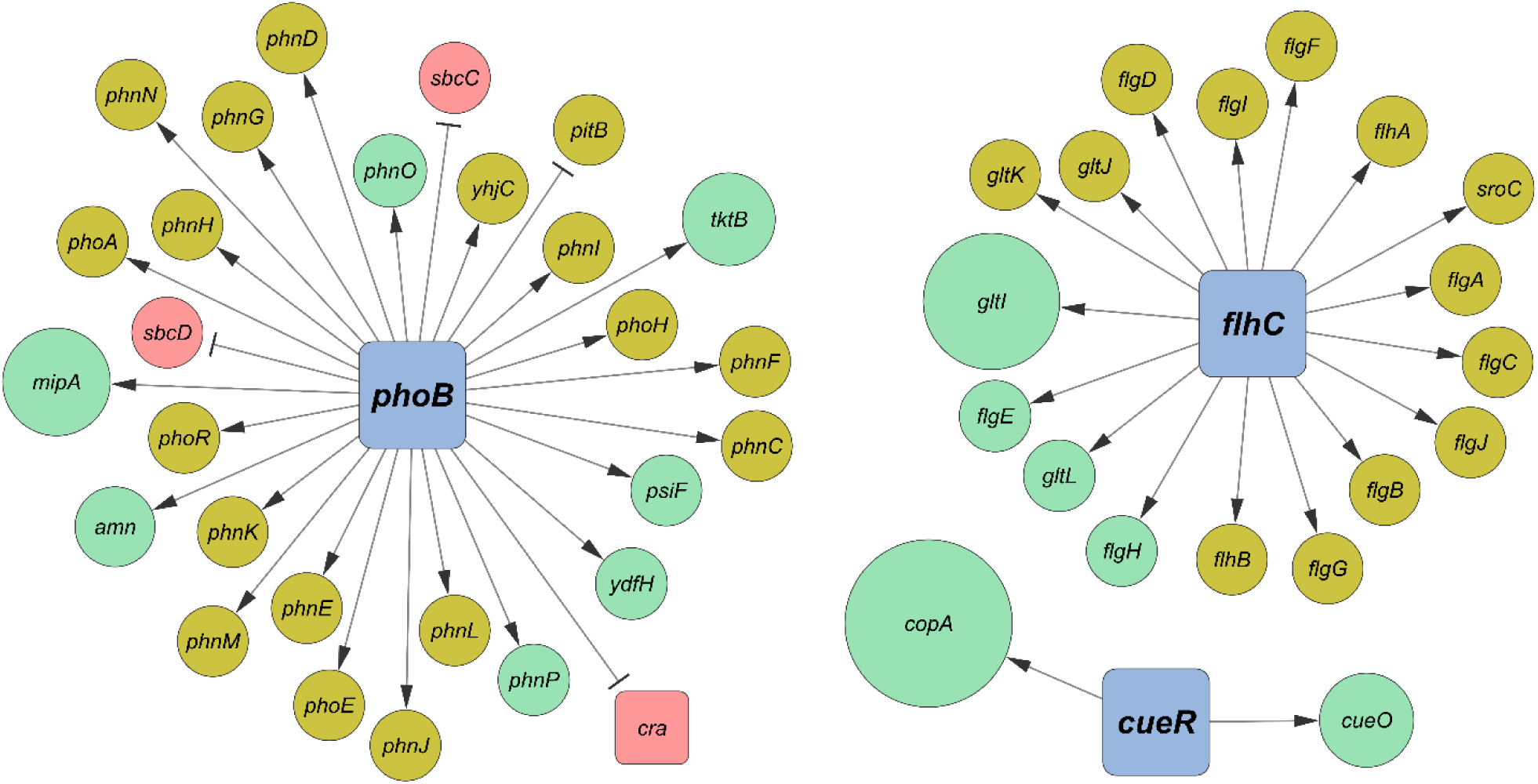
Regulatory subnetwork of predicted gene targets of the PFC mutant; green circles represent predicted silenced targets, red circles predicted induced targets and yellow circles genes with no proteomic coverage; size of the circles is proportional to the PL of the target.

**Supplementary Figure 6.**
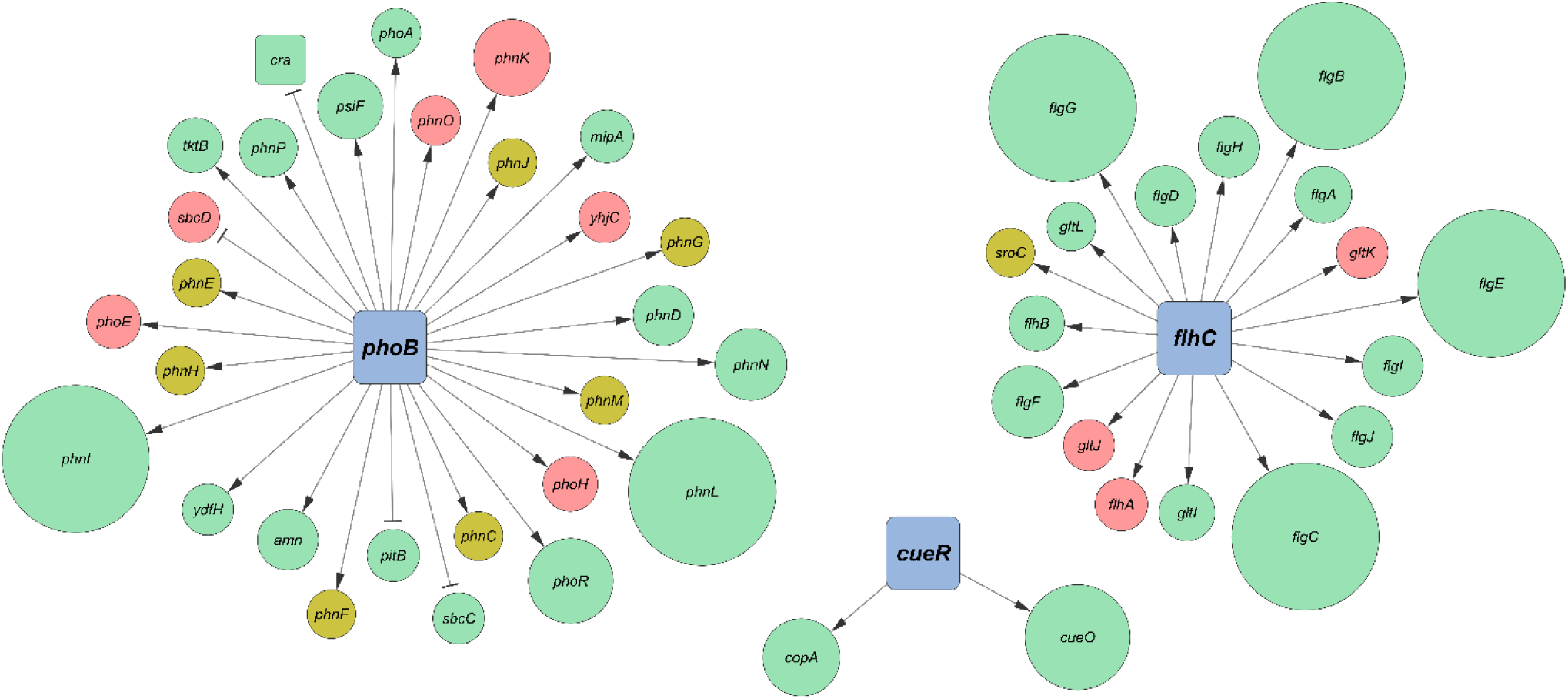
Integration of transcriptomics with computational tool predictions, the size of the circle corresponds to the fold change of each target (biggest circles represent fully silenced genes), in all cases green represent targets liberating resources (down regulated), red circles represent targets generating burden (up regulated) and yellow circles are targets not expressed.

**Supplementary Figure 7.**
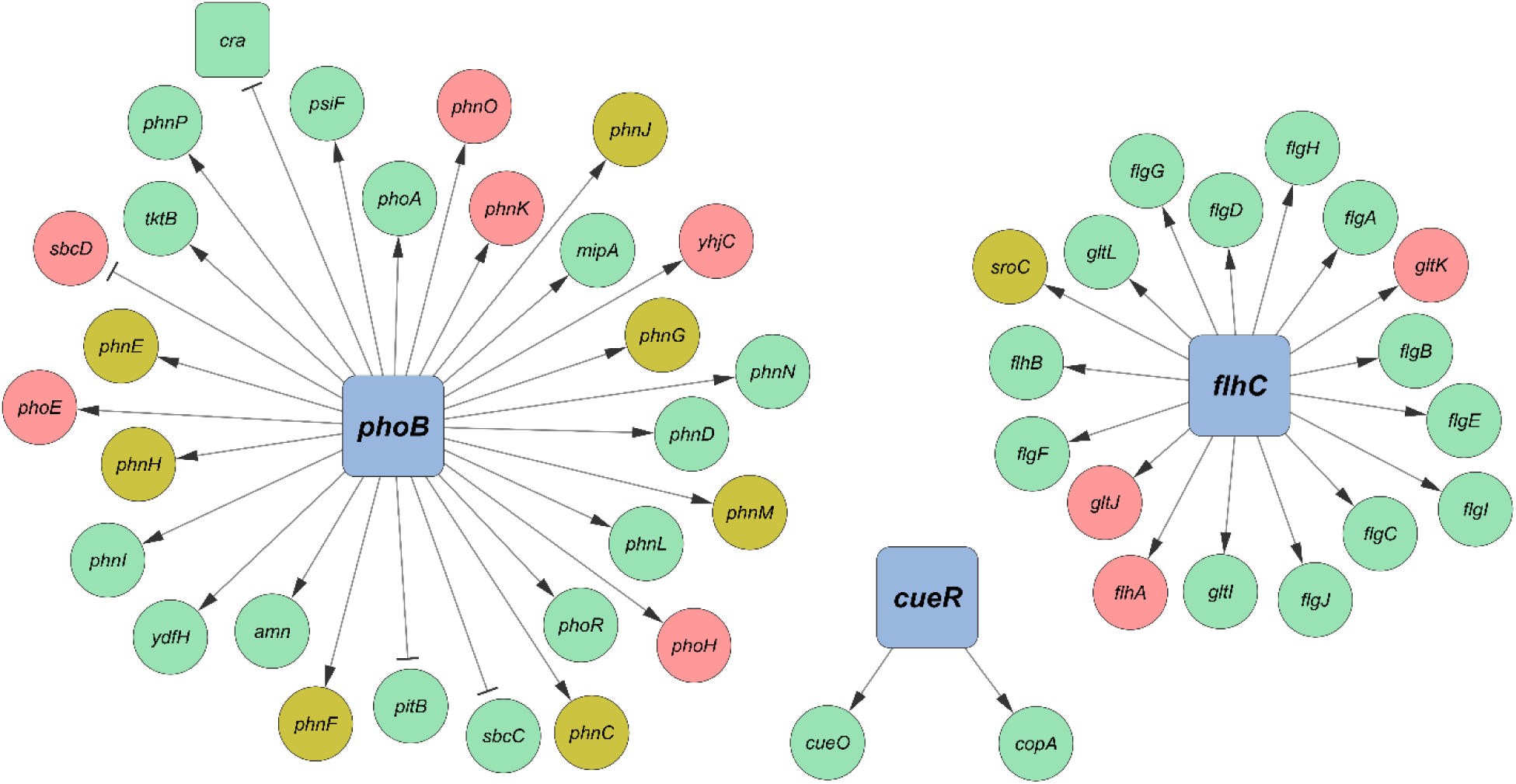
Accuracy of computational tool predictions (PFC) based on RNAseq data. Red circles represent wrong predictions, green circles represent accurate predictions and yellow circles represent unmapped predictions (expression was not detected).

**Supplementary Figure 8.**
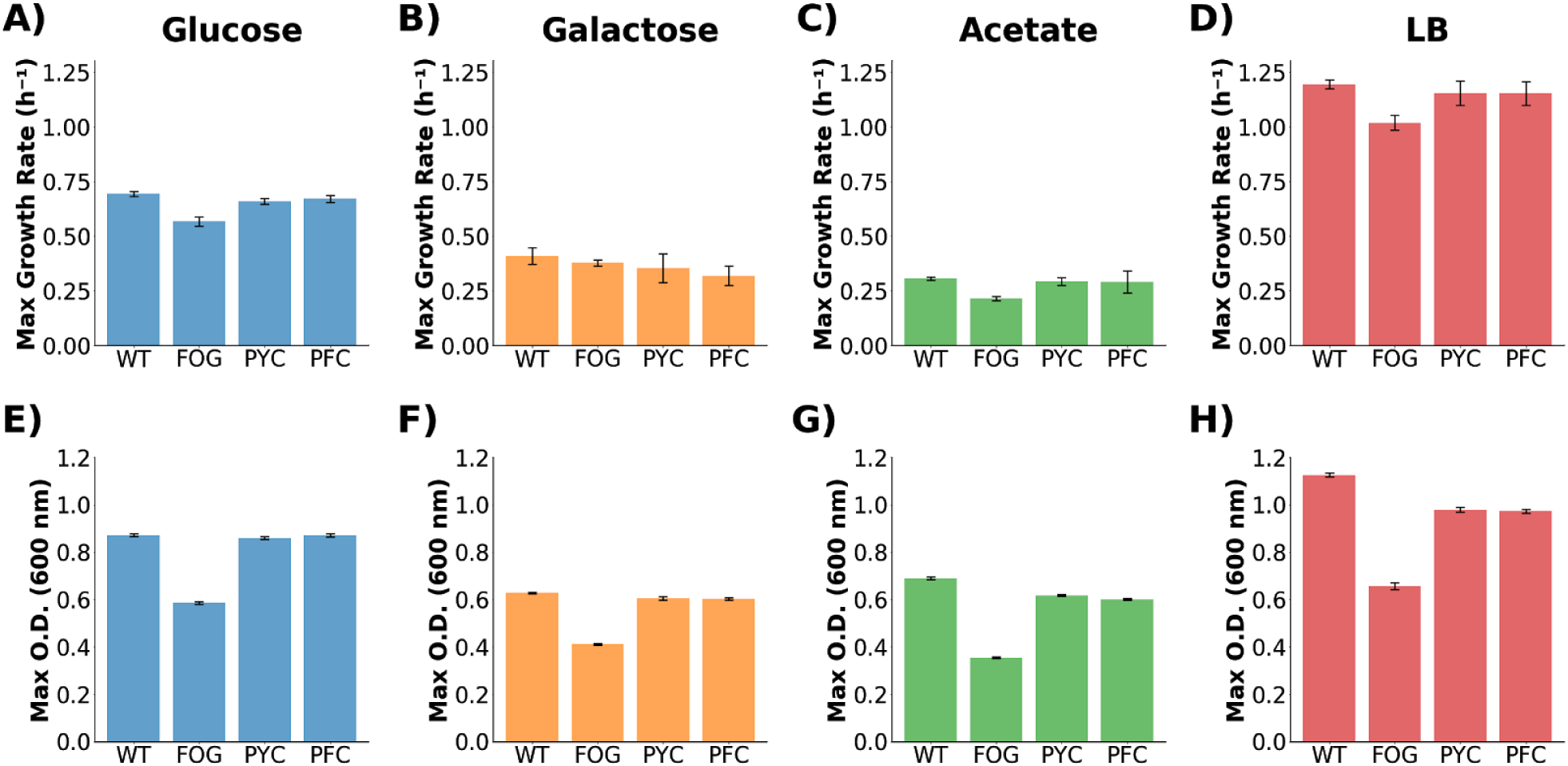
Phenotypic evaluation of strains on different carbon source supplemented M9 media and rich media (LB). (**A-D**) shows max growth rate and (**E-H**) shows max O.D.

**Supplementary Figure 9.**
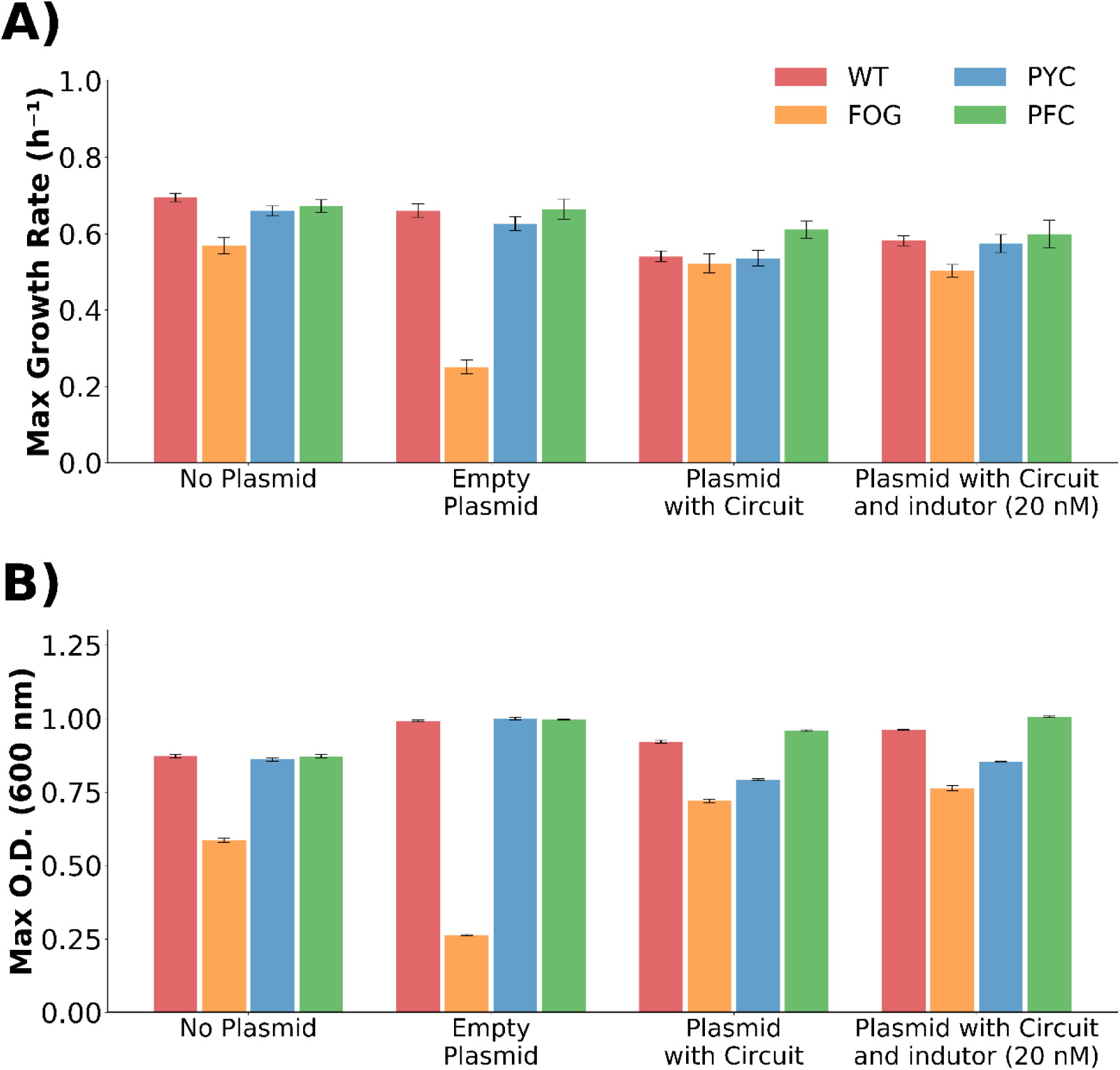
Metabolic burden while carrying empty, circuit plasmid and induced circuit plasmid, **A)** shows max growth rate and **B)** shows max O.D.

**Supplementary Figure 10.**
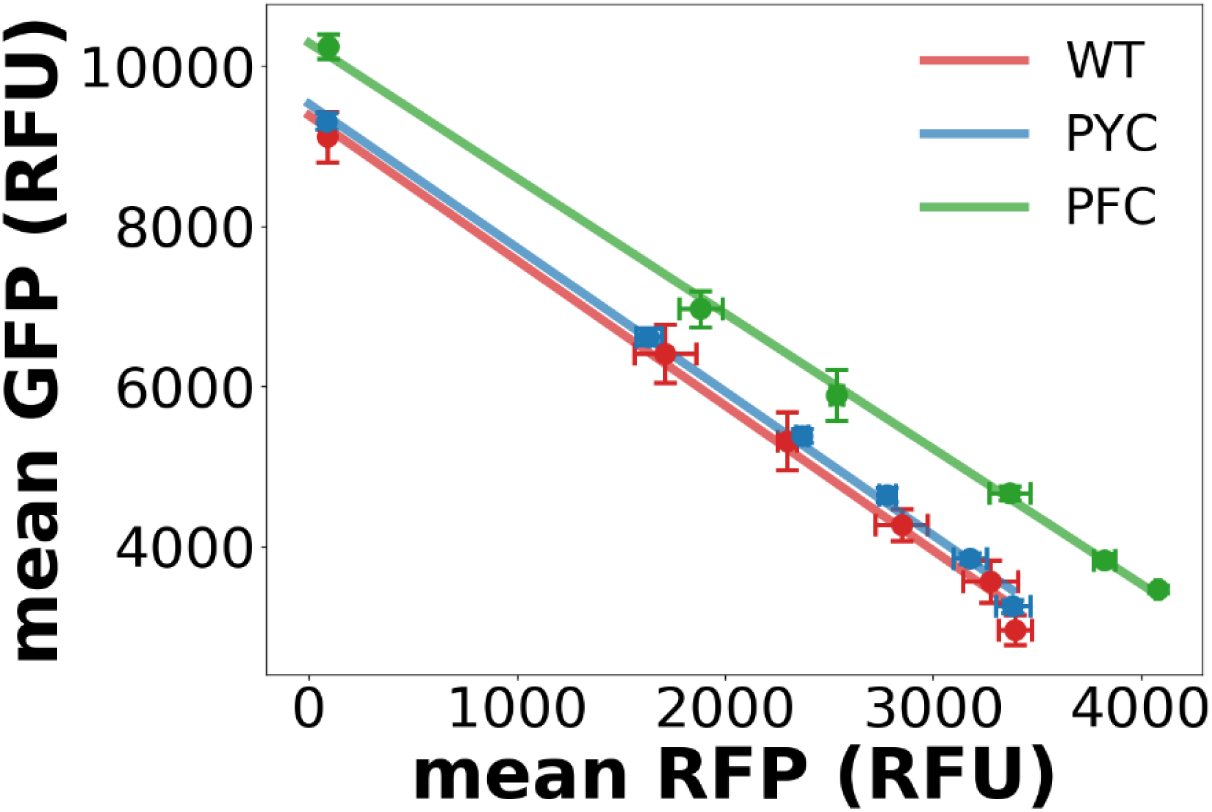
Isocost lines showing mean fluorescence per cell measured by flow cytometry during balanced growth (∼5 h).

**Supplementary Figure 11.**
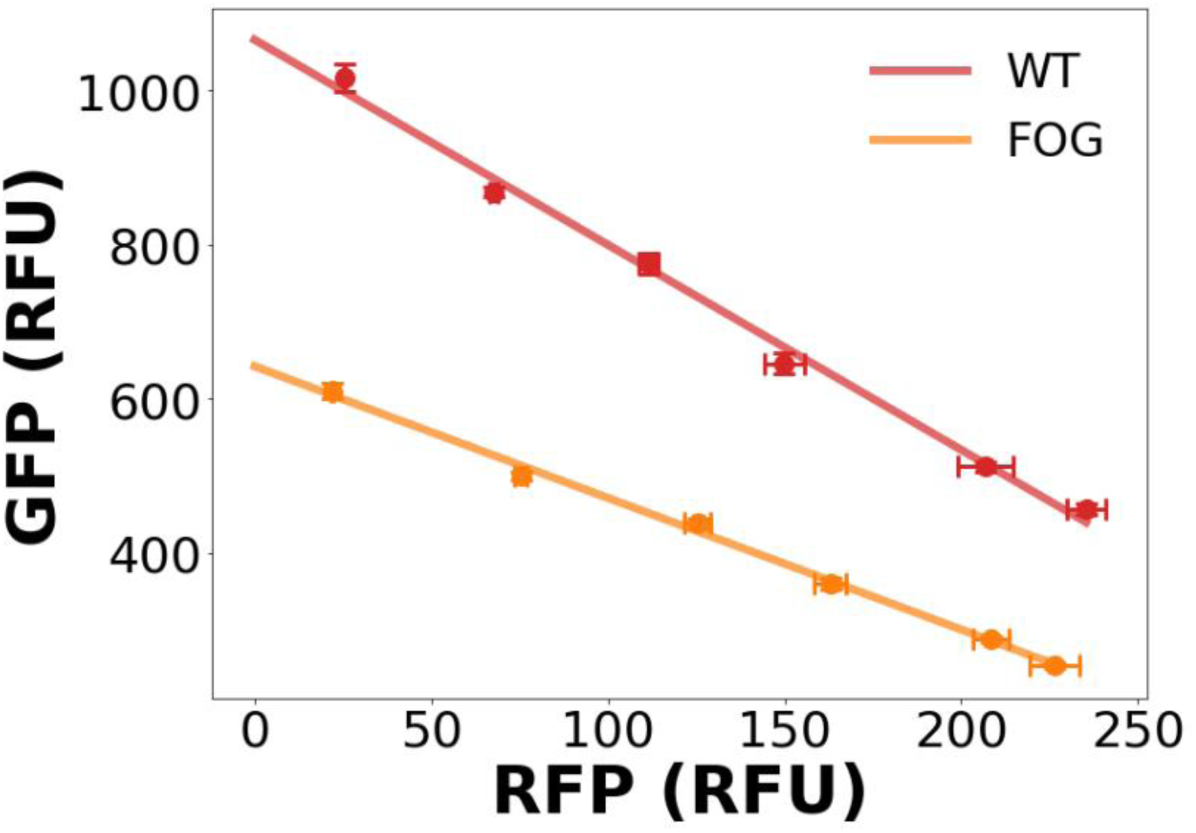
Isocost lines of the FOG mutant compared to the WT strain during balanced growth (∼5 h).

## Notes

#### Summary of Updates

It is the same text, just the abstract was divided in abstract and importance proper HTML tags

